# Potential clinical value and influence of conductivity in conductive cardiac patches for reducing post-MI arrythmia risks

**DOI:** 10.1101/2023.12.27.573394

**Authors:** Yuchen Miao, Zhenyin Fu, Juhong Zhang, Yuhang Tao, Kai Pang, Chengjun Wang, Qianqian Jiang, Liyin Shen, Tian Xia, Peixuan Lu, Zhen Xu, Ling Xia, Lijian Zuo, Jizhou Song, Changyou Gao, Dongdong Deng, Ruhong Jiang, Yang Zhu

## Abstract

Conductive cardiac patches can help to restore electric signal conduction of the diseased myocardium after myocardial infarction (MI). However, none of the conductive cardiac patches reported in literature has entered clinical trials. Bench-to-bedside translation of conductive patches has long been hindered by the lack of knowledge of the optimal patch conductivity and deep understanding of the potential clinical benefits and risks in patients. Here, we first prepared conductive cardiac patches with conductivities covering 5 orders of magnitude (10^-3^-10^1^ S/cm). Disagreeing with the mainstream opinion that patch conductivity close to native myocardium (10^-3^-10^-2^ S/cm) is most favorable, our results showed that patches with conductivity two orders of magnitudes higher than native myocardium (10^-1^-10^0^ S/cm) are most effective in restoring cardiac conduction and lowering inducibility quotient. Conduction velocity (CV) is the essence of the observed results. Rat experiments showed that the low-conductivity patch could increase the CV of infarcted myocardium, but did not fully compensate the mismatch in the CVs of infarct and healthy myocardium. Moderate-conductivity patches could increase myocardial CV to the same level of healthy myocardium, while high-conductivity patches further increased myocardial CV, causing a reversed mismatch. The relationship between patch conductivity and improved CVs in myocardium can be explained by monodomain model theory. Based on the theory, 3D finite element simulation of a MI patient heart predicted that a suitable, patch-improved myocardial CV could reduce the number of reentrants, and stabilize the remaining reentry circuits in the myocardium of the MI patient, which indicated its clinical value.

## Introduction

Cardiovascular disease is one of the main causes of global mortality, among which myocardial infarction (MI) has a high mortality rate [1]. After MI, cardiomyocytes (CMs) in the myocardium are unable to maintain normal excitation-contraction coupling process, and the electric signal propagation of the infarcted region is greatly hindered. The uneven electrophysiological distribution can lead to arrhythmias including ventricular tachycardia and ventricular fibrillation [2], causing angina pectoris and even cardiac arrest. Therefore, rebuilding electrical conduction in the MI regions is important in myocardial repair. Many studies have reported that conductive biomaterials improved electrical signal conduction in infarcted myocardium [3-5]. Conductive biomaterials are able to transmit electromechanical, electrochemical, and electrical stimuli to cells [6], thus have the potential to synchronize remaining CMs in infarct with adjacent myocardium [7]. In addition, electrical stimulation through conductive biomaterials can effectively regulate maturation and functionalization of CMs, which could be useful when cell therapy is combined [8].

Conductive biomaterials used for myocardial repair mainly include intrinsic conductive polymers, carbon-based nanomaterials, and metal nanomaterials [9]. Generally, conductive fillers are mixed in non-conductive polymer scaffold or hydrogel to prepare conductive cardiac patches or injectable hydrogels. However, conductive cardiac materials constructed by such method have low conductivity (10^-7^-10^-2^ S/cm) [10] due to poor continuity of the internal conductive network, and have a small conductivity span range (only 2-5 times) due to the limitation of conductive fillers concentration. The majorities of researchers [11-13] believe that the moderate conductivity of the material is similar to that of the myocardium, but lacks theoretical or experimental evidence. At the same time, detailed and thorough studies are needed to reveal the essence of conductive patch restoring MI electrical conduction [14, 15].

Different from the strategies of fabricating conductive biomaterials for cardiac repair, 3D connected graphene oxide (rGO) aerogels are employed as the conductive substrate, and methacrylated whey protein (WPI-MA) hydrogel was used to fill the conductive network of rGO aerogels, providing mechanical support. Through this method, five types of patches with conductivity of 10^-3^ S/cm, 10^-2^ S/cm, 10^-1^ S/cm, 10^0^ S/cm, 10^1^ S/cm were successfully prepared, having a wide conductivity span. Real-time electrocardiogram and extended MI electrocardiographic induction experiment have demonstrated that the key to improve electrical conduction in infarcted myocardium is to increase conduction velocity (CV) of the electrical signals in the myocardium by the patches. At the same time, simulation on the hearts of MI patients have shown that reentrants in the hearts can be decreased in a patch conductivity dependent manner, as a result of patch-enhanced CV. We also explained and predicted the therapeutic effects with the monodomain model theory.

## Materials and methods

### 1. Fabrication of the conductive patches

Aqueous GO suspension (8 mg/ml, Hangzhou Gaoxi Technology, China) was cast-dried on a polyethylene glycol terephthalate (PET) substrate to obtain a GO film. The GO films were immersed into a hydrazine hydrate aqueous solution (N_2_H_4_, Sinopharm Chemical Reagent, China) for 60 min at room temperature to fabricate GO aerogels. During the foaming process, the N_2_H_4_ solution could preliminarily reduce GO. By controlling the GO aerogel reduction conditions (Table 1), rGO aerogels with different conductivities were prepared.

**Table 1.**
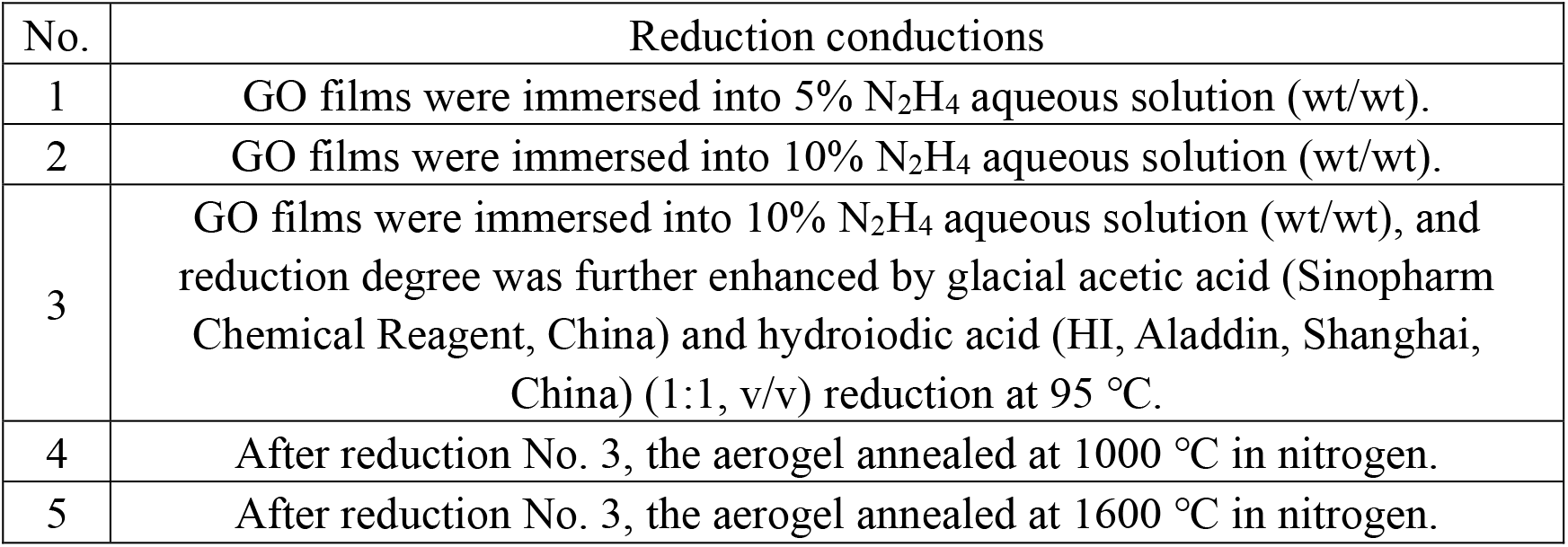
The reduction conductions for GO aerogels.

| No. | Reduction conductions |
| --- | --- |
| 1 | GO films were immersed into 5% $\text{N}_2\text{H}_4$ aqueous solution (wt/wt). |
| 2 | GO films were immersed into 10% $\text{N}_2\text{H}_4$ aqueous solution (wt/wt). |
| 3 | GO films were immersed into 10% $\text{N}_2\text{H}_4$ aqueous solution (wt/wt), and reduction degree was further enhanced by glacial acetic acid (Sinopharm Chemical Reagent, China) and hydroiodic acid (HI, Aladdin, Shanghai, China) (1:1, v/v) reduction at 95 °C. |
| 4 | After reduction No. 3, the aerogel annealed at 1000 °C in nitrogen. |
| 5 | After reduction No. 3, the aerogel annealed at 1600 °C in nitrogen. |

WPI powder was purchased from Tianjin Milkyway Import & Export, China. According to our previous work [16], 10 g WPIs were dissolved at room temperature in 190 mL of 1X PBS solution, magnetically stirred for 20 minutes. Subsequently, 2 mL of methyl methacrylate was added, and the mixture pH was adjusted to 8 by 2 M sodium hydroxide and stirred for 24 hours at room temperature. The final solution was dialyzed in ultrapure water for 3 days using a 1000 Mw dialysis bag. The dialyzed solution was frozen at -20 °C for one day and then freeze-dried for 48 hours to obtain WPI-MA powder.

rGO aerogels were immersed into methacrylated WPI (WPI-MA) solution (15 wt% in 1X PBS, with 2% ammonium persulphate). Through an insulin needle, the WPI-MA solution was injected into the rGO aerogels along its periphery until fully fill the aerogels. Subsequently, residue air was completely removed by vacuum, and WPI-MA solution air pressured into the rGO aerogels. This step was cycled 3 to 5 times. The filled aerogels were heated at 65 °C for 30 minutes to crosslink WPI-MA in rGO aerogels, resulting in a composite of rGO aerogels and WPI-MA hydrogel, which is the conductive cardiac patch.

### 2. Conductivity measurement of conductive patches

The internal pore structures of rGO aerogels and conductive patches were observed via field emission scanning electron microscope (SEM, S-4800, Hitachi, Japan).

Conductivity of patches were calculated from I-V curves (measured by 2400 SourceMeter, Keithley, with four-electrode method) by the formula:

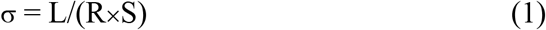

where σ is electrical conductivity, L is electrode distance, R is resistivity, and S is the cross-sectional area of patches. LED light bulbs were used to display patch conductivity changes on dynamic hearts.

### 3. Patch implantation in rats

Male SD rats (8 weeks, 200-250 g), purchased from Zhejiang Academy of Medical Sciences, were randomly divided into 7 groups (n=10, for each group): sham group, MI group, patch groups (5 groups corresponding to 5 different conductivities). All animal experiments were approved by the Institutional Animal Care and Use Committee of Zhejiang Experimental Animal Center (ZJCLA-IACUC-20010275). The MI rat model was established as described in previous studies[17]. After rats anesthetized with 1% pentobarbital, the left anterior descending (LAD) artery was ligated with 6-0 silk sutures in MI group. Rats in the sham group was only subjected to thoracotomy treatments. In patch groups, patches (circular, 8 mm diameter) were sutured onto rat hearts.

### 4. Fabrication of flexible sensor array

The preparation process of a flexible sensor array was as follows: first, the polyimide (PI) prepolymer was spun coated and high-temperature cured to obtain a polyimide film with a thickness of 15 μm. The sensor units and interconnecting wires were fabricated on the surface of the PI film by electron beam evaporation and photolithography method. The array was arranged into 6 columns and 5 rows based on the electrocardiographic sensor units. By dry etching, the PI between sensor units and outside the circuit was removed to obtain a hollow flexible sensor array, which ensured contact between the myocardium and the conductive patch. The anisotropic conductive film (ACF) cable was joined to the contact pads of the sensor array through hot pressing bonding, and the other end was joined to the SAGA (TMSI, The Netherlands) instrument to achieve electrocardiogram signal acquisition. The TMSI Polymer Data Manager software recorded and analyzed the signals.

### 5. Real-time electrocardiograms (ECGs)

The flexible sensor array was directly attached to the rat epicardium to record real-time ECGs of rats in all groups. ECGs of healthy rat were recorded first, then the same rats underwent MI modeling surgery, and the ECGs were recorded 30min post MI. The rat hearts were treated with the 5 types of conductive patches with different conductivities. ECGs with and after removing the patches were recorded in all patch groups.

### 6. Electrical stimulation

The ECGs test adopted the Leads I connection method, and the three electrodes were connected to the iWorx data acquisition system containing IROX-RA-384 (iWorx) and iWrie-BIO8 (iWorx). ECGs were recorded by LabScribe software in the electrical stimulation process. At 4 weeks after implantation of patches, arrhythmias were induced by an external electric stimulation catheter (FTS-1913A-1018, iWorx). The catheter was passed through the internal carotid artery to the left ventricle. Programmed electrophysiological stimulation (PES) includes burst pacing, S1S2, and S1S2S3. When there were 15 or more rapid ventricular rhythms, the rats were undergone sustained ventricular arrhythmias, indicating successful induction[18]. If arrythmia was not induced after the PES protocol, 5 mg/mL of isoproterenol at a dose of 0.1 mL/100 g would be injected. The induction was performed again according to the PES protocol, and the inducibility quotient was calculated. According to the statistical method of rank sum test, the difficulty of successfully inducing arrythmia was scored.

### 7. Simulation of influence of patches on MI patient hearts

Simulation on one myocardial infarction (MI) patient from Beijing Anzhen Hospital was performed (approval by the Institutional Review Board of Beijing Anzhen Hospital). Cardiovascular magnetic resonance-late gadolinium enhancement (CMR-LGE) images of this patient were collected and to construct the heart models. Cardiac MRI were acquired by 3.0 T scanner (Sonata, Siemens, Erlangen, Germany) or GE scanner (DISCOVERY MR 750w; GE, Boston, MA, United States). The detailed image acquisition protocol can be found in the previous published literature [19].

All analyses and measurements related to the simulation model were performed using custom software developed in MATLAB (MathworksInc., USA). Experienced experts manually segmented the epicardial and endocardial boundaries of each two-dimensional slice from the LGE images. The pixels between these boundaries were considered as myocardium. Subsequently, the modified Gaussian mixture model method was employed to automatically identify the infarcted regions[20]. Finally, the full width at half max method was used to further segment the infarct tissue into gray zone and core scar. The detail process could be referred to our previously published paper [20].

After Image segmentation, the myocardial tissue was evenly divided into ten layers using an algorithm based on the Laplace-Dirichlet rule. The outermost part was designated as ‘patch/tissue interface’ (its thickness was small and did not significantly affect the tissue model underneath it), and it was simulated as passive tissue with conductor-like properties (conductivity corresponds to different electrical signal conducting capacities). The model was paced from 19 ventricular sites, including 17 sites on the LV (left ventricular), 1 near the right ventricular outflow tract, and 1 at the right ventricle apex, according to American Heart Association (AHA) Classification Standards[21]. By recording the reentry points at the 19 sites, the therapeutic effects of different patches on arrhythmia were analyzed.

### 8. Statistics

All data were presented as the means ± standard deviation and analyzed using Graphpad Prism. Two-tailed student’s t-test was employed to determine whether two groups differed significantly. Multiple groups were tested by one-way analysis of variance (ANOVA) with Tukey post-hoc method. P-value of < 0.05 was considered statistically significant. *p<0.05, **p<0.01, ***p<0.001, ****p<0.0001.

## Results

### 1. Fabrication and characterization of the conductive patch

As shown in Fig. 1A, GO aerogels were obtained through foaming of GO film. N_2_H_4_ released nitrogen, hydrogen, and ammonia gas, which infiltrated the layers of GO film, spread the layers to form a highly porous structure (porosity>99% ). At the same time, N_2_H_4_ preliminary reduced GO aerogels. Subsequently, chemical reduction and thermal reduction were carried out to obtain rGO aerogels with higher conductivities. In order to stabilize aerogels geometry and hence the conductivity, WPI solution was injected into rGO aerogels and crosslinked in-situ. SEM images show that GO layers were separated and formed a continuous network structure with average pore diameter 56±0.5 um (Fig. 1B). WPI hydrogel completely fill the pores of rGO aerogels without destroying its network structure.

**Figure 1.**
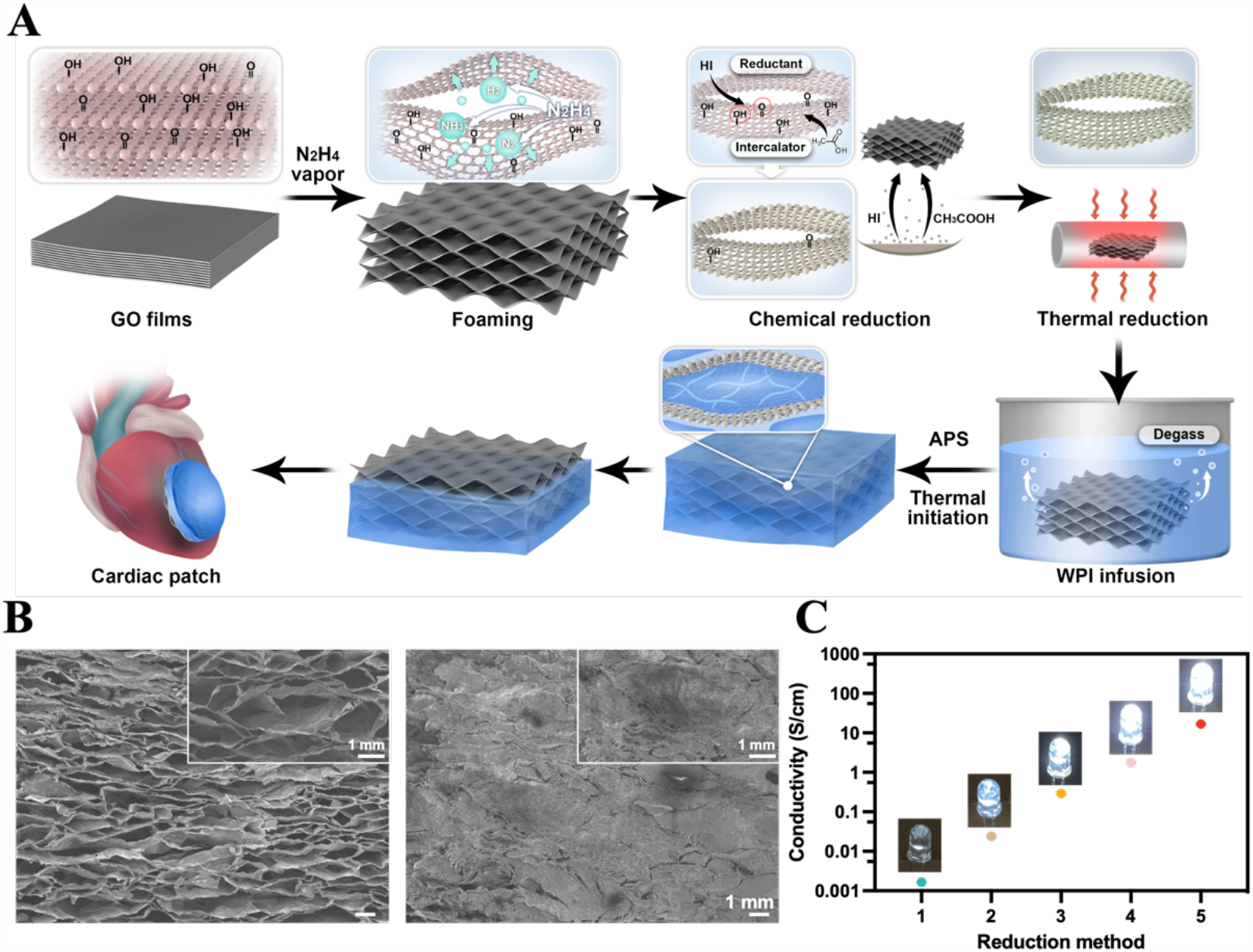
Preparation and characterizations of the conductive cardiac path. (A) The preparation process of the conductive patch. (B) The internal structures of rGO aerogel and the conductive patch. (C) Patch conductivity and corresponding brightness changes of LED light bulbs.

The conductivities of rGO aerogels fabricated with the five different reduction methods were measured to be 1.65×10^-3^ S/cm (method 1), 2.39×10^-2^ S/cm (method 2), 2.93×10^-1^ S/cm (method 3), 1.76×10^0^ S/cm (method 4), 1.70×10^1^ S/cm (method 5), respectively (Fig. 1C). The difference in conductivity between adjacent two points was about 10 times, indicating the successful preparation of conductive patches with conductivity ranging from 10^-3^ S/m to 10^1^ S/cm, covering 5 orders of magnitude. The brightness of LED bulbs under same voltage can displayed the conductivity increase with the increase in reduction degree.

### 2. Conductive patches improved electrical signal conduction and reduced arrhythmia

The real-time ECGs was detected by a customized flexible sensor array to evaluate the immediate therapeutic effect of the conductive patch on MI rats. The sensor was attached to rat epicardium. Attributed to the hollow structure of the sensor array, the conductive patches can directly contact the cardiac tissue (Fig. 2A). ECGs were recorded before, during and after patch contact with infarcted hearts.

**Figure 2.**
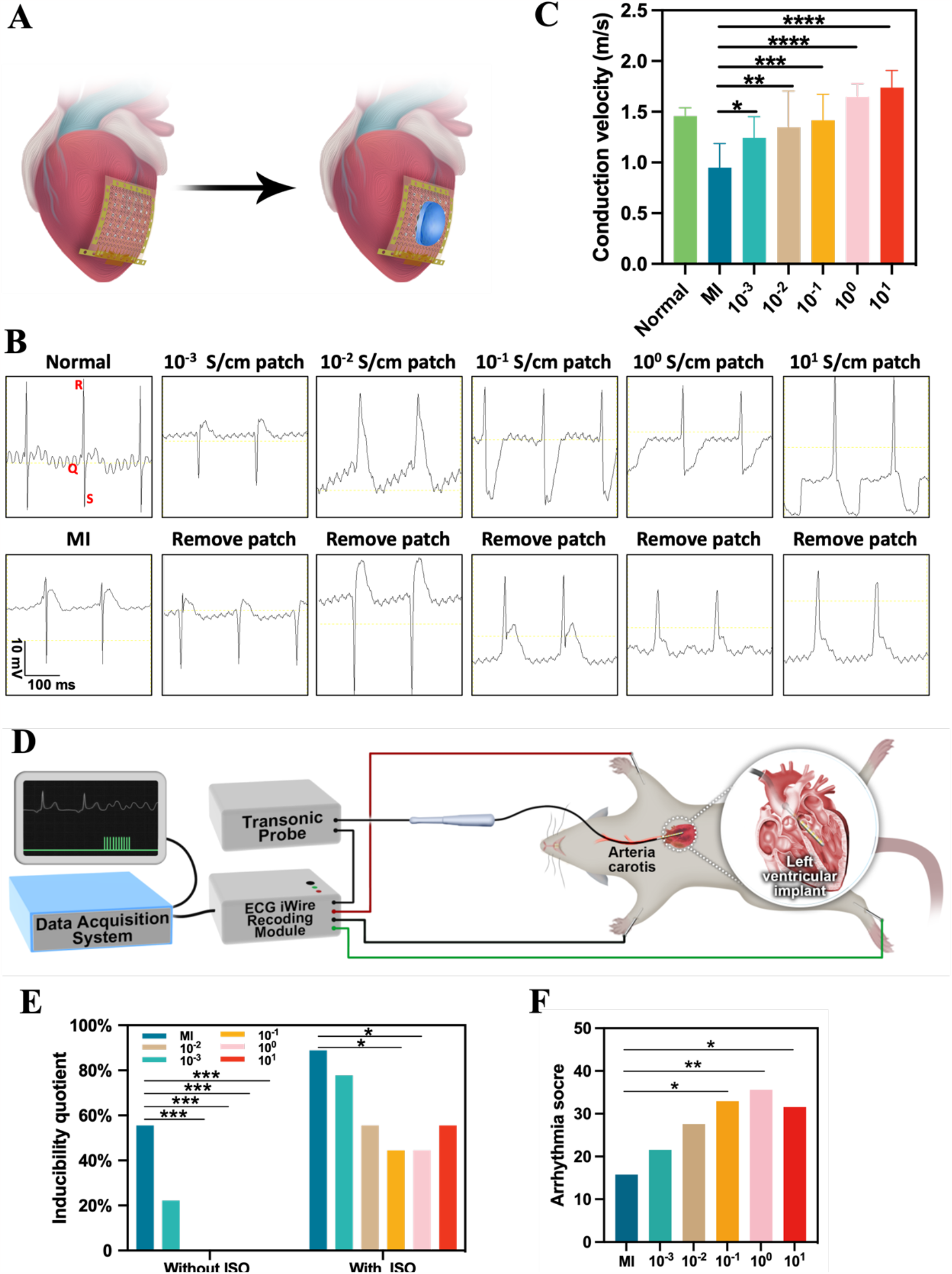
Electrophysiological repair of myocardial infarction with different conductivity patches. (A) Scheme of real-time ECGs experiment, (B) ECGs changes before and after patch treatment (C) corresponding CV of different patches. (D) Scheme of induction experiment, (E) the inducibility quotient, (F) the arrhythmia score.

The QRS complex of healthy rats was relatively narrow. The flexible sensor obtained the same results as it was attached to rat chest. In infarcted myocardium, the depolarization vector disappears or weakens, altering the QRS complex vector loop, and resulting in “erosion” on the vector. The post-MI ECGs showed typical features, including QRS complex widening, ST segment elevation, Q wave deepening, and notches (Fig. 2B). Compare to the ECGs before patch treatment, the 10^-3^ S/cm patch cleared the notches. The 10^-2^ S/cm patch narrowed the QRS complex, depressed the ST segment, and eliminated the Q wave deepening, but the QRS internal was still longer than that of healthy group. The ECGs, after 10^-1^-10^1^ S/cm patch treatment, generally recovered to the healthy state, but their ST segment was depressed. Clinically, ST segment depression occurs in endocardial damage, while ST segment elevation occurs in epicardial and transmural damage. After patch treatment, the ECGs changed from ST segment elevation to depression, which may be due to the patch rebuilding the electrical conduction in the epicardium, possibly in certain aspects more electrically active compared to healthy epicardium.

Based on the changes in the peak positions of QRS complexes from the 30 ECG channels collected by the sensor array, conduction velocity (CV) in the infarct region corresponding to each group was calculated (Fig. 2C). The measured CV of healthy myocardium is 1.46±0.08 m/s, post-MI CV decreased to 0.95±0.24 m/s. Compared to MI rats, CV of myocardium covered by patches with conductivities of 10^-3^-10^-1^ S/cm increased by 30.5%, 42.1% and 49.5% and were recovered to 1.24±0.21 m/s, 1.35±0.36 m/s and 1.42±0.26 m/s, respectively. CV of myocardium covered by patches with conductivities of 10^0^-10^1^ S/cm were all faster than that of healthy myocardium, which were 1.65±0.13 m/s and 1.74±0.0.17 m/s, respectively.

In addition to real-time ECGs, we also induced arrythmia 4 weeks after patch implantation in rats via programmed electrical and chemical stimuli, to evaluate the capability of conductive patches in reducing the risks of post-MI arrythmia in a more clinical relevant setting (Fig. 2D). Electrical stimulation alone (without isoproterenol (ISO)) only induced ventricular tachycardia in MI rats and the 10^-3^ S/cm patch treated rats. With both electrical stimulation and ISO injection, all groups were induced arrythmia. Among the patch groups, the 10^-1^-10^0^ S/cm patch groups had the lowest inducibility quotient of 44%, significantly smaller than that of the MI group. By assigning scores to the induction status of each group through rank sum test, the two groups with the highest scores were the 10^-1^-10^0^ S/cm patch groups, indicating that these two groups of patches have the best inhibitory effect on arrhythmia.

### 3. Appropriate conduction velocity can reduce reentrant

On the personalized 3D modeling of an MI patient’s heart, 19 sites were selected for reentrant stimulation according to AHA Classification Standards. Reentrant can be divided into persistent and non-persistent reentrant. The persistent reentrant is further divided into anatomical and functional reentrant. Non-persistent reentrant refers to that the number of reentrant loops generated at the lesion location is less than three, and it does not pose a threat to patients. Persistent reentry has three or more reentrant loops. Anatomical reentry typically occurs in the isthmus between the semi-infarcted area and the infarcted tissue. The reentry loop is obvious and has a fixed turn back route. Functional reentry is less stable than anatomical reentry, but the position of the reentry loop does not change much. As shown in Table 2, when there was no patch, a large number of reentrants were generated. After patch implantation, the reentrant in the original stimulus points almost disappeared, but new reentrant was generated at other stimulus points. As the conductivity increased, the number of reentrant showed a “decrease, then increase” trend, which was similar to the results of arrythmia induction experiments. When there was no patch, there were more anatomical reentrants compared to functional ones, but with the addition of patches, the number of anatomical reentrants was significantly reduced, that may be attributed to the improvement in electrical conduction between the infarcted and semi-infarcted dead zones by patches.

**Table 2.** The reentrant of MI and different conductivity patches under ventricular tachycardia.

| Conductivity (S/cm) |  |  |  |  |  |  |  |  |  |
| --- | --- | --- | --- | --- | --- | --- | --- | --- | --- |
| MI | | $10^{-2}$ | | $10^{-1}$ | | $10^0$ | | $10^1$ | |
| Site | Reentrant type | Site | Reentrant type | Site | Reentrant type | Site | Reentrant type | Site | Reentrant type |
| 9 | A | 1 | A | 7 | F | 6 | F | 1 | F |
| 10 | A | 6 | A |  |  | 9 | A | 6 | F |
| 11 | A | 7 | A |  |  | 18 | F |  |  |
| 12 | A | 10 | F |  |  |  |  |  |  |
| 13 | A |  |  |  |  |  |  |  |  |
| 14 | F |  |  |  |  |  |  |  |  |
| 15 | F |  |  |  |  |  |  |  |  |
| 19 | A |  |  |  |  |  |  |  |  |
| Reentrant num. |  |  |  |  |  |  |  |  |  |
| 8 |  | 4 |  | 1 |  | 3 |  | 2 |  |

Tips: A=Anatomical reentrant, F=Functional reentrant.

### 4. The essence of different therapeutic effects is conduction velocities

CV in LV wall (across the infarct) was measured (Table 3), patches with conductivity similar to that of healthy myocardium (10^-3^-10^-2^ S/cm) did not improve the CV of the infarcted heart to healthy level, while patches with conductivity 10 to 1000 times higher than that of the myocardium (10^-1^-10^1^ S/cm) significantly increased CV and even surpassed the healthy heart. Corresponding changes were also observed on the real-time ECGs. The low-conductivity patches (10^-3^-10^-2^ S/cm) did not effectively shorten the QRS interval, while the 10^-1^-10^1^ S/cm patches were able to shorten the QRS interval, to a level similar to the healthy state (Fig. 2B). At the same time, the increase in CV also reduced the probability of arrhythmia under programmed electrical and chemical stimuli, as shown in Fig. 2E. The 10^-1^-10^0^ S/cm group had the lowest inducibility quotient (0% without ISO, 44% with ISO), while when the CV was too high (1.74±0.17 m/s, the10^1^ S/cm group), the risk of arrhythmia increased compared to those of the 10^-1^-10^0^ S/cm groups, probably due to the mismatch of the CV in high-conductivity patch treated myocardium and adjacent normal myocardium as the CV in latter was relatively slow.

**Table 3.** CV calculated through experiments and simulated by monodomain model.

| Conductivity (S/cm) | $10^{-3}$ | $10^{-2}$ | $10^{-1}$ | $10^0$ | $10^1$ |
| --- | --- | --- | --- | --- | --- |
| CV of experiments (m/s) | $1.24 \pm 0.21$ | $1.35 \pm 0.35$ | $1.42 \pm 0.26$ | $1.65 \pm 0.13$ | $1.74 \pm 0.17$ |
| CV of monodomain model (m/s) | 0.205 | 0.771 | 1.084 | 1.112 | 1.135 |

In order to further validate the results of small animal experiments, we also simulated MI patient with different conductivity patches. Based on the reentrant mechanism, the mechanism of different conductivity patches producing different number of reentrant points could be explained (Fig. 3). After MI, the CV in the infarcted area was decreased, forming a slow-fast conduction mismatch with the surrounding normal myocardium, which increases the probability of reentrant. The low-conductivity patches could improve the CV to a certain extent, so the probability of reentrant is reduced. The moderate-conductivity patch could further increase the CV to that of the surrounding normal myocardium, thus making it difficult to induce ventricular tachycardia. The high-conductivity patches, due to their fast CV, generated a reversed mismatch in CVs of the infarcted area and surrounding normal myocardium, in which conduction of electrical signals was faster in patch-covered infarcted area, resulting in new reentrants.

**Figure 3.**
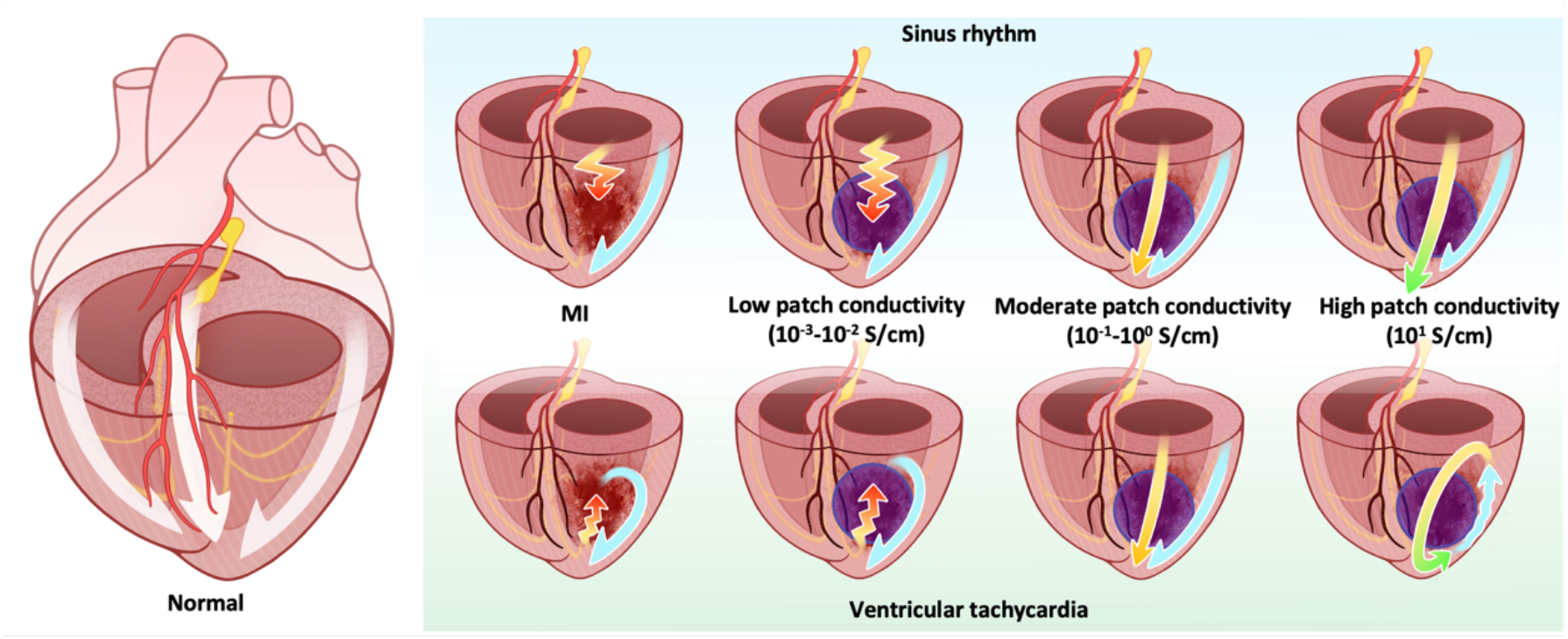
The explanation of the reentrant mechanism of patches with different conductivities.

To explain why patches with different conductivity leads to different CV in the treated myocardium, we first tested the RC circuit hypothesis. It is assumed that healthy myocardium on both ends of the infarct area form a capacitor, and the patch served as a resistor connecting the two halves of the capacitor. The infarct between the two halves of the capacitor is considered as an insulator, no current going into, through or out the patches. The time required for upper stream activated cells to excite downstream resting cells can be calculated as follows:

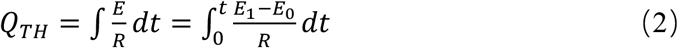

Q_TH_ is the threshold charge for generating excitation, E_0_ is the membrane potential of cardiomyocytes (CMs) in an excited state, E_1_ is the membrane potential of CMs in a resting state, R is the resistance of the patch, combined with formula (1) and the CV calculation formula V=L/t (V is the conduction velocity, L is the length of the patch, and t is the conduction time of the electrical signal on the patch). The relationship between the CV and corresponding conductivity is obtained as 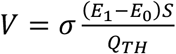, indicating their linear relationship, but it was clearly inconsistent with the actual experimental results (Table 3).

To match the measured results and model predicted results, the diffusion of electrical excitation in myocardial tissue is described with a monodomain model (simplified dual domain model) [22]. The monodomain model divides myocardial tissue into two regions: intracellular and extracellular, with equal anisotropy. Based on the assumption of the monodomain model, the propagation equation of CMs action potential are as followed:

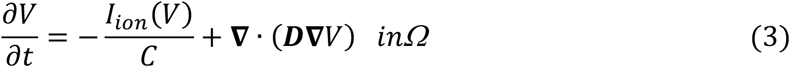

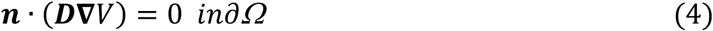

Where ∂v/ ∂t is the time derivative of the transmembrane potential, I_ion_ is the total ion current, C is the battery capacitance per unit surface area, and D is the diffusion tensor. Ω and ∂Ω represent the domain of interest and its boundaries, respectively, and **n** is an outward unit vector perpendicular to ∂Ω [22]. Table 3 showed the myocardial CV corresponding to patches with different conductivities, calculated through experiments and monodomain model. Although the simulated CV was slightly lower than the corresponding experimentally measured CV treated by the same patches, the trend of CV changes with the increase of conductivity was the same as the experimental ones, which proved that monodomain model theory could semi-quantitatively explain the experimental results, particularly how patches with different conductivity improves CV and the differences in the resulting myocardial CVs.

## Discussion

The repair in electrical conduction system cannot be explained by a simple RC circuit composed of myocardial tissue and patches, but rather by the theory of combining the RC circuit composed of patches with the monodomain model of myocardial tissue. This is in line with the characteristics of the native myocardial electrical conduction system, so that it can predict the CV of myocardium under different conductivity patch coverage. To date, theoretical basis of conductive biomaterial improving electrical signal conduction of infarcted myocardium is qualitative. A widely accepted and typical explanation is that patches can bridge the healthy myocardium surrounding the infarct by converting the ionic conductivity of the myocardium into the electronic conductivity of the patch [23], but with no quantitative explanation why patches can improve CV of infarcted myocardium.

It is widely believed and accepted that in terms of improving cardiac conduction, biomaterial conductivity is better when it is close to healthy myocardium, ∼10^-3^ S/cm. The conductivity of patches used in this study spans five orders of magnitude (10^-3^-10^1^ S/cm) and can maintain a stable conductivity on beating hearts, which is desirable in isolating the contribution of conductivity. In previous studies, there was no material with such a large conductivity range, and no demonstration that material conductivity can remain stable even during heart beating. The correspondingly designed flexible sensor used in this work can directly measure the myocardial electrical signal (ECG) covered by the patch, which reflects the real-time treatment effect of the patch on the infarcted myocardium and supports calculation of CV at each point before and after patch treatment. Due to technical limitations, the previous conductive patch works were unable to directly measure the myocardial conduction under patch coverage. In addition, this article also conducts induction experiments, which have more clinical significance and can reveal risks that ECGs cannot detect. There are only 5% previous work reporting arrythmia induction experiments.

The 10^-1^-10^0^ S/cm patches group can instantly shorten the QT interval of MI rats and increase the CV of the myocardium to slightly higher than that of healthy myocardium. With increased CV, the probability of arrhythmia is minimized, the probability of reentrant is also minimized. So, the most suitable patch conductivity for MI treatment should be 100 to 1000 times higher than that of healthy myocardium.

## Limitations

The main conclusions of this study is derived from rat experiments, but there are significant differences between infarcted rat hearts and patient hearts. Infarct area in rat MI model is more continuous, but that of MI patients is often scattered in healthy myocardium. Therefore, the quantitative relationships obtained from the rat model may need to be modified to match patient conditions. Particularly, reentrant results is obtained via simulation and is not supported by experimental data. Finally, because the conductive patches were stitched onto the heart, thus there may be inconsistency in patch/myocardium contact, which might increase the variation in therapeutic effects.

## Conclusion

Our study showed that increasing the electrical signal conduction velocity in infarcted myocardium is the essence of repairing the electrical activities in MI damaged hearts by conductive biomaterials. The optimal patch conductivity is 100-1000 times higher than that of healthy myocardium, which lead to improved CVs slightly higher than that of healthy myocardium. By increasing CV, patches with suitable conductivities can recover ECG pattern to close to normal instantly after contacting the hearts. Correspondingly, inducibility quotient of arrhythmia and the number of reentrant were all decreased. However, when patch enhanced CV in infarct is too high, it increases the risk of arrhythmia and generates new reentry points. Therefore, a patch with appropriate conductivity can not only restore the electrical connection in the infarct area, but also reduce the number of reentrant points, which could be helpful for clinical radiofrequency ablation surgery.

## Acknowledgements

This study was financially supported by the National Key Research and Development Program of China (No. 2019YFE0117400), National Natural Science Foundation of: China (No. 82202328), the Starry Night Science Fund of Zhejiang University Shanghai Institute for Advanced Study (No. SN-ZJU-SIAS-004).

## Notes

### Competing Interest Statement

The authors have declared no competing interest.

